# EEG correlates of perceived food product similarity in a cross-modal taste-visual task

**DOI:** 10.1101/814939

**Authors:** M. Domracheva, S. Kulikova

## Abstract

The most common tools to understand perception of food products are hall-tests, surveys, observations, etc. However to get reliable results these approaches require large samples, making them costly and time-consuming. Furthermore, they are also highly expert-dependent and rely on the assumption that study participants can express their preferences consciously and explicitly. Here we suggested an EEG-based approach to evaluate perceived product similarity in a cross-modal taste-visual task. Two candidate neurometrics measured from Fz electrode were tested: the amplitude of N430-620 from evoked response potentials (ERP) and the power of induced gamma oscillations during 400-600 ms period after visual stimulus presentation. Both suggested metrics showed a strong correlation with the perceived similarity scores at both individual and group levels, however N430-630 had greater inter-subject variability making it less suitable for practical applications. The results based on the power of induced gamma oscillations (N=18) not only reflected the results from traditional hall-tests (N=200) but also allowed better discrimination between different food products.

## Introduction

Nowadays, healthy eating trends are substantially influencing the choice of consumers. According to Nielsen research (2015, 2016) up to 70-80% of the world population make their food choices taking into account various healthy food attributes such as organic origin (absence of GMOs, artificial flavorings and colorants), low levels of cholesterol, salt, sugar and fat, etc. Following this trend, food manufacturers, including sweet producers, are trying to include in their product lines items that have these healthy attributes (or at least are believed to have them). This often leads to the emergence of new innovative products made from unconventional ingredients or/and using innovative technologies.

When a manufacturer launches such new products he/she faces questions regarding product positioning and potential consumers’ perception of the product. The most common tools used in traditional marketing to understand consumer perception of food products are hall-tests, surveys, observations, interviews and focus groups. However these approaches require a large sample to be used in order to ensure the reliability of the results, making such investigations costly (Wright, 2005). At the same time, data collection and processing using traditional marketing methods are also highly prone to researcher’s subjectivity (Wright, 2005), leading to low results reliability. Furthermore, previous studies have shown that quite often consumers are not able to express their preferences consciously and explicitly (Stasi et al., 2018), suggesting why results from such studies may fail to explain real consumer behaviour.

Neuromarketing approaches, which aim at recording and interpreting product-relevant brain activity using neurophysiological techniques like EEG (electroencephalography) or fMRI (functional magnetic resonance imaging), may help to overcome these limitations and provide traditional marketing techniques with new complementary information on true consumers’ preferences (Ariely, Berns, 2010; Vlăsceanu, 2014). However, to our knowledge, there is no proposed neuromarker that could be used to assess perceived similarity between innovative and familiar food products. Thus, the goal of the present study was to suggest a neuromarketing approach that may be reliably applied for investigation of perceived similarity between different food products on relatively small sample sizes.

## Materials and Methods

### Test product

Multi-cereal candies (MCC) with creamy and chocolate flavours were considered as test products because our preliminary analysis of the customers feedback has revealed that this product is perceived rather ambiguously: some consumers indeed perceive it as a candy, others as cookies, granola, cereal min-bars or waffles. Although, generally, MCC are believed to be healthier than regular sweets or candies, it is often noted that they are nevertheless very rich in calories (200-400 kcal/100 g) and may not be suitable for maintaining fitness. Such ambiguous perception of MCC made them an excellent candidate for development of a neuromarketing strategy that would evaluated subjectively perceived similarity between different food products. We investigated two different flavours (creamy and chocolate) as we hypothesized that adding cacao to MCC would shift its perception towards less healthy products and make it more candy-like.

### Hall test

The first step of this study was conducted in a form of traditional hall tests in order to get preliminary hypothesis about different food categories that consumers may associate with MCCs. Hall tests were carried in two formats on two different samples: in a “Blind” hall test respondents (N=99; 34 males, 65 females, mean age = 20.8±3.3) were testing products with their eyes closed being unaware of the testing product, while in the “Open Eyes” (N=101; 29 males, 72 females, mean age = 21.2±2.5) hall test they could see the product and its package. In addition to MCC we also tested a piece of boiled broccoli and a piece of fried potatoes. Both of these products were extremely unlikely to be associated with MCC, with broccoli being perceived as an example of healthy food and fried potatoes as unhealthy. Using both “Blind” and “Open Eyes” conditions allowed, on the one hand, to evaluate the perception of the MCC taste characteristics, excluding the influence of other factors as much as possible, and on the other hand, to evaluate the potential effect of packaging on MCC perception, since it may happen that the used packaging could shift MCC perception towards its perceiving as a candy.

After testing each product (MCC, broccoli or fried potatoes), participants were asked to compare using 5-points Likert scale the tested product with 8 other products: candies, waffles, cookies, broccoli, fried potatoes, muesli/granola, cereal bars and oatmeal. Next, they were asked to indicate using 10-points Likert scale to what degree the tested product and each of the products from the list above could be considered as a healthy food product. Finally, they were asked about how much they liked the tested product and answered the questions from the Health and Taste Attitude Scale (HTAS) (Roininen et al., 2001), which measures the importance of perceived aspects of utility and taste in the food selection process. All data collected at this stage will be available at demand from the corresponding author upon acceptance of this study for publication.

### EEG recordings

The second step of our study aimed at identifying novel neuromarkers that could be used to evaluate perceived similarity between different food products. Experiments were performed in 18 healthy volunteers (mean age=24.7±3.6) and each participant signed an informed consent before the experimental session. Similar to the first stage of our study, half of the experiments was conducted under a “Blind” condition, and the other half – under “Open Eyes” condition. EEG was recorded using a 24-channel wireless EEG system (Neurotech, Taganrog, Russia) with a standard 10/20 electrode positioning.

Each experimental session included 8 blocks. At the beginning of each block, participants tasted one MCC. After having it tasted and after the disappearance of artifacts related to chewing and swallowing, a random sequence of 40 images of various products was shown on the computer screen. This sequence included 5 different images for each of the 8 product types considered at the first stage of the present study. Each image was presented for 3 sec. After each image, participants evaluated using a 5-points Likert scale how much the tested product matched the product presented in the image (from 1 – “these are completely different products” to 5 – "these are exactly the same products”). The time for response was not limited. In between the blocks, participants could rest as long as they wished and water was provided *ad libitum*. At the end of the experimental session participants were asked about how much they liked the tested product and completed the HTAS questionnaire.

### Data analysis

EEG data was filtered in 0.1-100 Hz range and two different neurometrics were extracted: 1) power of induced gamma oscillations (30-80Hz) between 400 and 600 ms. after each image presentation normalized by the power of ongoing gamma oscillations during the 200ms period preceding stimulus presentation; and 2) an amplitude of negative difference component (N430-620) in the evoked response potentials (ERP).

Comparisons of paired distributions (obtained from the same sample under one experimental condition) of perceived product similarities and healthiness was performed using Wilcoxon signed-rank tests. Comparison of unpaired distributions (different samples from “Blind” and “Open Eyes” sessions) were performed using Mann-Whitney tests. Relationships between perceived product similarities and EEG metrics were evaluated using Spearman correlation.

## Results

### Hall test

The results of the hall test have shown that participants equally liked both creamy and chocolate MCC. During the “Open Eyes” testing the mean liking scores were 3.96 and 3.92 correspondingly (p>0,82). However, under “Blind” condition testing the liking scores tended to be higher (p=0,082) for both types of MCCs with creamy flavour (mean liking score=4.42) prefered (p<0,021) to chocolate one (mean liking score =4.06).

Under “Open Eyes” conditions creamy MCCs were mainly associated with cereal bars (mean similarity score=4.23), while chocolate MCCs were perceived more as candies, cereal bars and granola (corresponding mean similarity scores were 3.65, 3.59, 3.31) (Fig.1). Under “Blind” conditions creamy MCCs were associated with waffles, granola and cereal bars (mean similarity scores = 3.72, 3.62, 3.54 correspondingly) and chocolate MMCs – with cereal bars and granola (corresponding mean similarity scores were 3.78 and 3.49).

**Fig.1.**
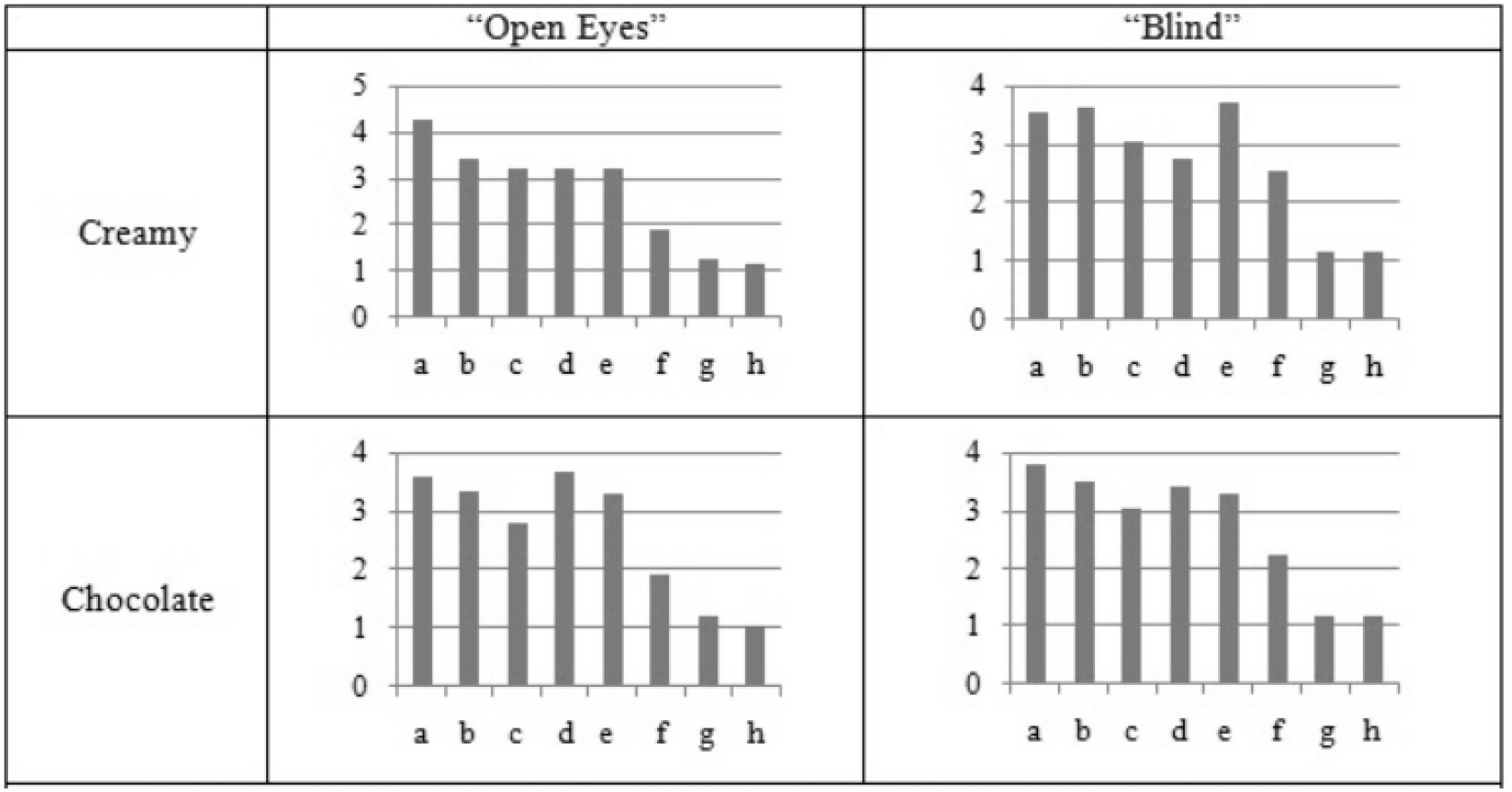
Perceived similarity for MCCs with other food products tested for two different flavours (creamy and chocolate) under two experimental conditions (“Open Eyes” and “Blind”). Notations: a – cereal bar, b – granola, c – cookies, d – candies, e – waffles, f- oatmeal, g – fried potatoes, h – broccoli.

Analysis of perceived product healthiness suggests that under “Open Eyes” conditions both types of MCCs were perceived healthier than fried potatoes (p<0.00001), candies (p<0.00001), waffles (p<0.00001) and cookies (p<0.00001), less healthy than broccoli (p<0.00001) and oatmeal (p<0.00001) and could be compared to cereal bars (p>0.5/0.2 for creamy/chocolate MCC correspondingly) or, to a bit lesser extent, to granola (p>0.04/0.1) or (Fig.2). Similar results were observed under “Blind” condition: both types of MCCs were considered healthier than fried potatoes (p<0.00001), candies (p<0.00001), waffles (p<0.00001) and cookies (p<0.00001) and less healthy than broccoli (p<0,001 / 0.00001) and oatmeal (p<0.00001). Creamy MCCs could be compared to granola (p>0,24) or cereal bars (p>0,045), however chocolate MCCs were considered less healthy than granola or cereal bars (p<0,01).

**Fig.2.**
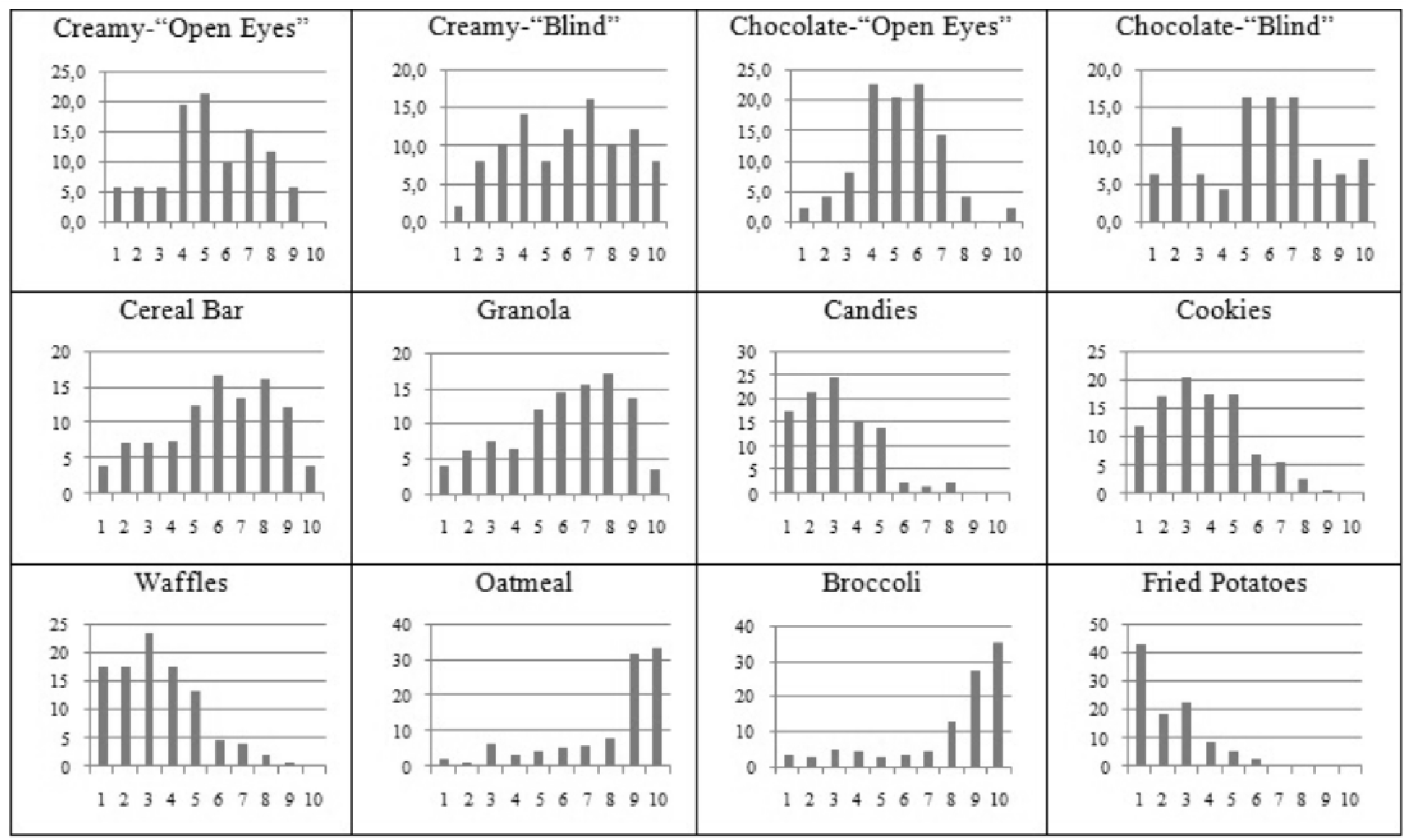
Distributions of perceived healthiness for two types of MCCs tested under “Open Eyes” and “Blind” conditions and for other investigated food products. Axis Y indicates the percentage of respondents that has put the corresponding score on a 10-point Likert scale for a given product.

### Comparison of Hall Test results and EEG metrics

Presentation of congruent images consistently evoked an increase of induced gamma oscillations at frontal Fz electrode during 400-600ms period after stimulus presentation (Fig.3), while incongruent images evoked an increase in N430-620 amplitude (Fig.4).

**Fig.3.**
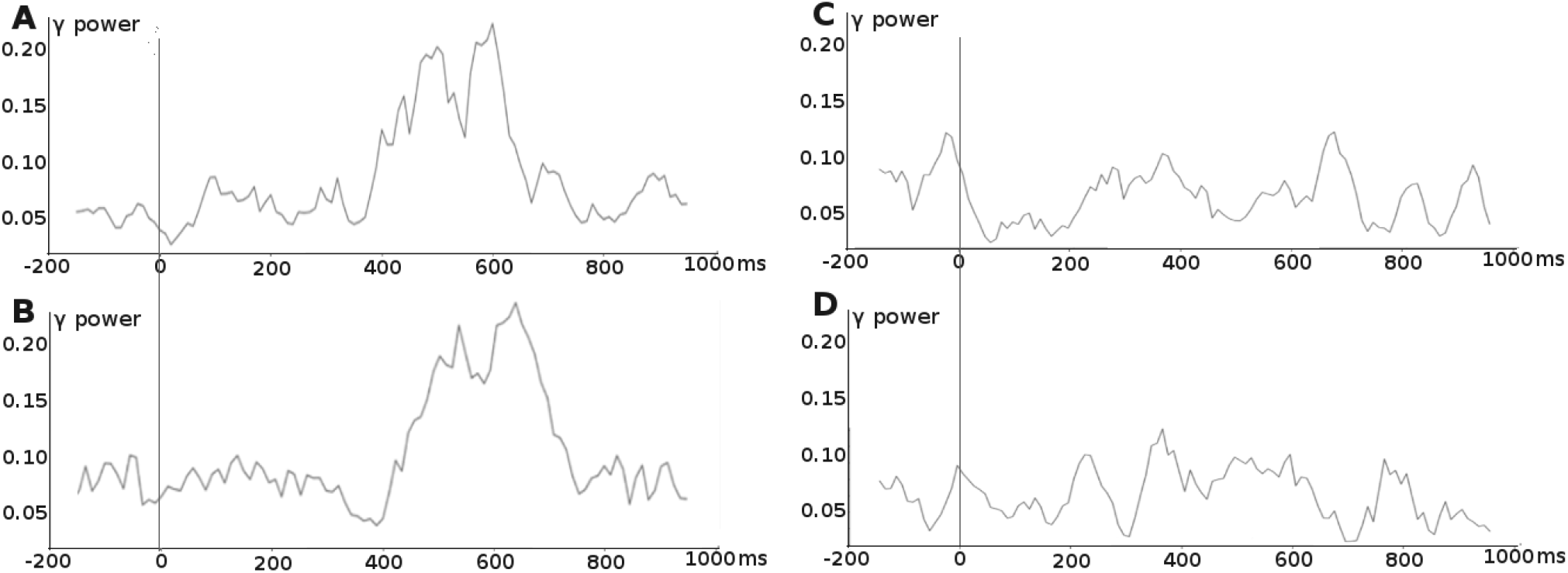
Representative examples of graphs showing dynamics of gamma oscillations on Fz electrode following presentation of congruent (A – cereal bar, B – granola) and incongruent (C – fried potatoes, D – broccoli) images. Stimulus was presented at 0-time point.

**Fig.4.**
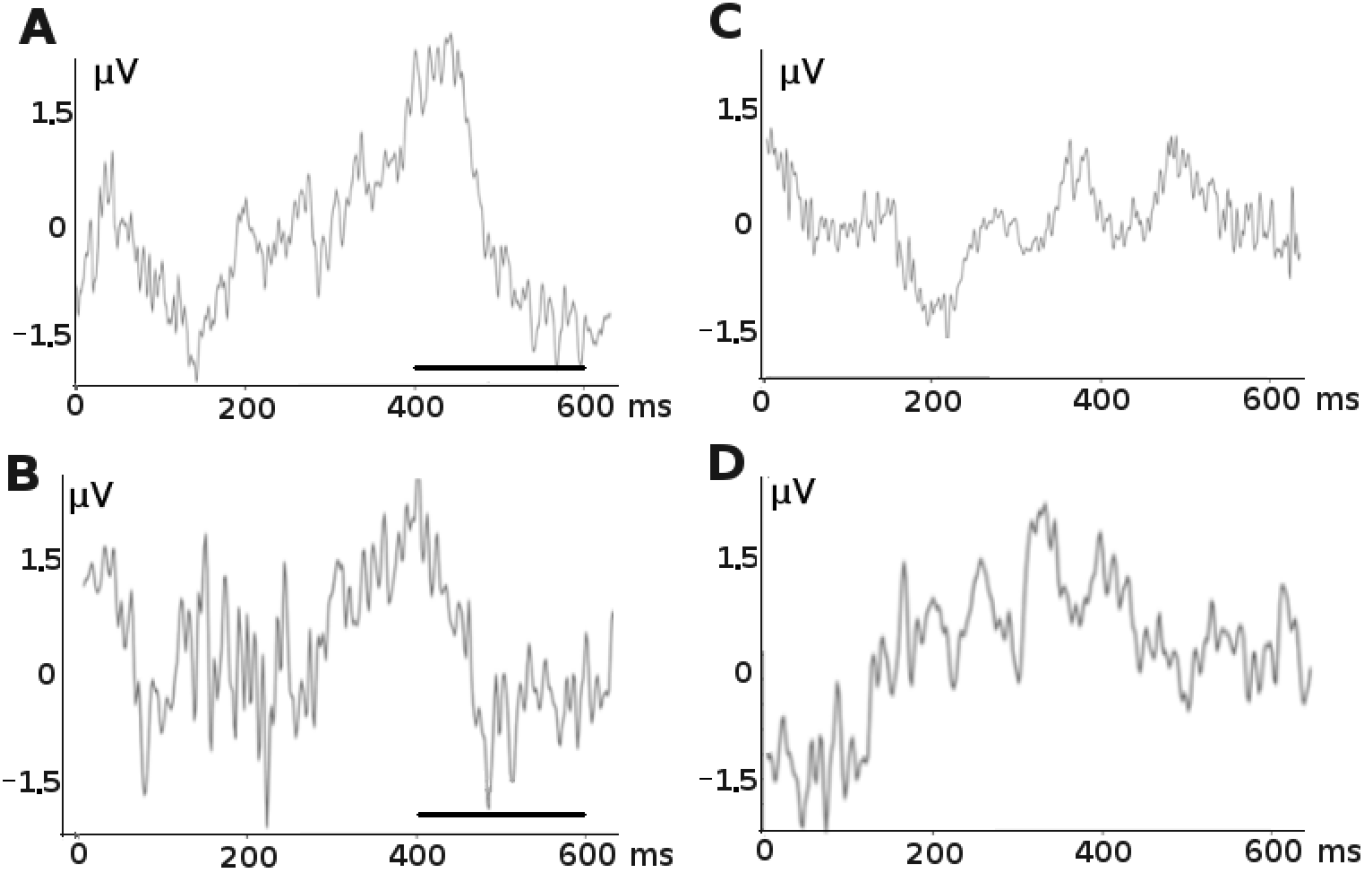
Representative EEG recordings at Fz electrode following presentation of incongruent (A – fried potatoes, B – broccoli) and congruent (C – cereal bar, D – granola) images. Stimulus was presented at 0-time point. Black bar indicated the presence of N430-630 component in recordings A and B.

Comparison of perceived product similarity and the two considered neurometrics revealed strong correlations between them both at individual and group levels (Fig.5,6). The perceived product similarity positively correlated with the power of induced gamma oscillations (Fig.5) and negatively with the N430-620 amplitude (Fig.6).

**Fig.5.**
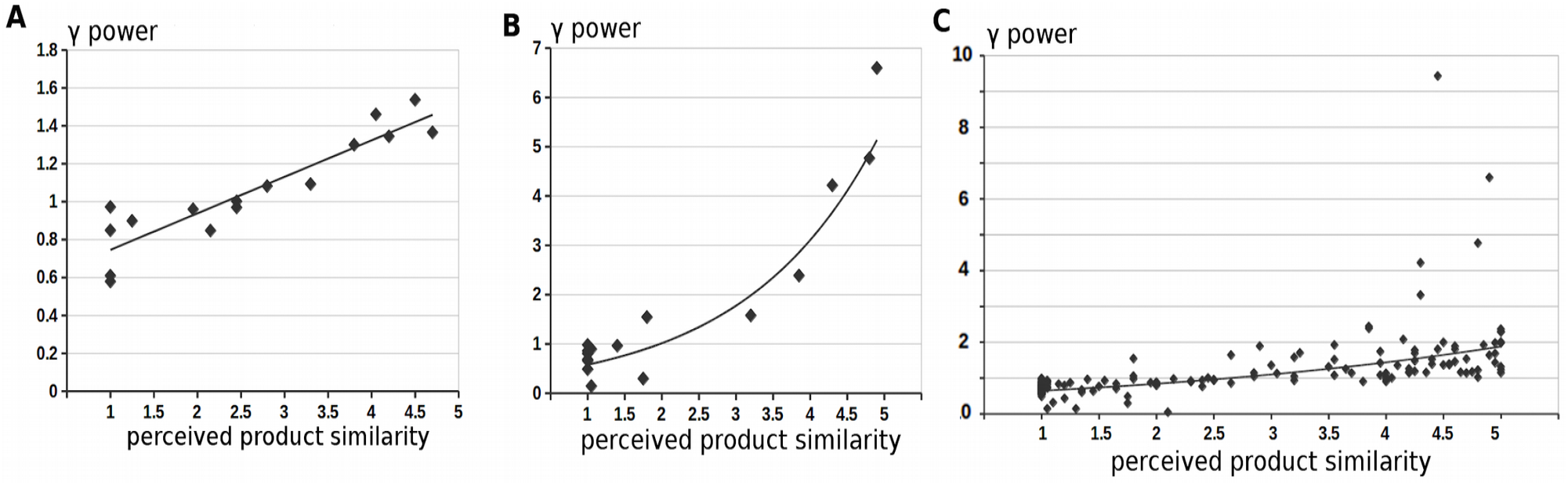
Graphs showing relationships between induced gamma power and perceived product similarity in individual participants (A, B) and on a group level (C, data from 18 participants). In the majority of cases the relationship was linear (A, R2=0.83; C, R2=0.69), whoever in two cases (for example, B) it was better explained by exponential curves (R2=0.81).

**Fig.6.**
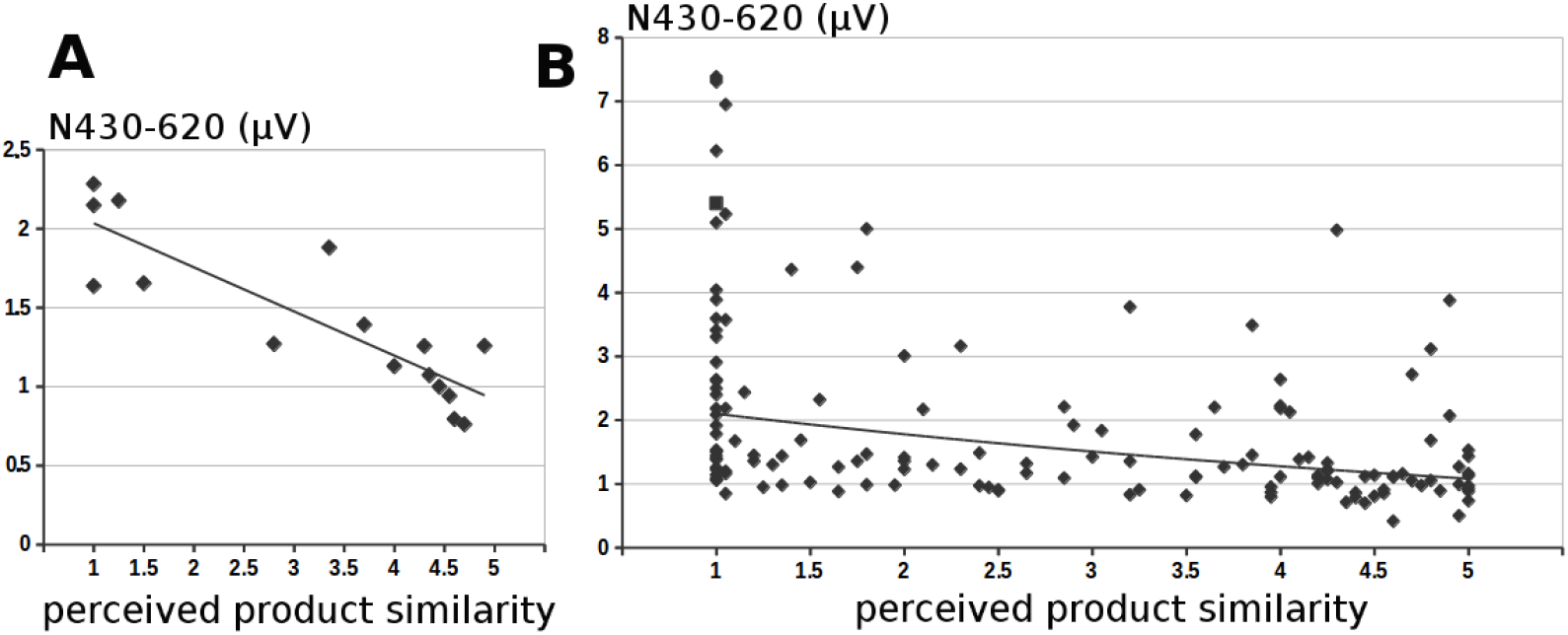
Graphs showing relationships between induced N430-620 amplitude and perceived product similarity in an individual participants (A, R2=0.74) and on a group level (B, data from 18 participants, R2=0.24).

For most of the subjects these relationships were linear, however in a couple of cases their were better explained by exponential curves (Fig.5B). Of importance, at the group level, the relationship between perceived product similarity and the power of induced gamma oscillations was much stronger than with N430-620 amplitude due to its large intersubject variability.

## Discussion

In this work we considered two potential neuromarkers to measure perceived similarity of food products: the amplitude of N430-620 from ERP and the power of induced gamma oscillations during 400-600 ms period after stimulus presentation. The choice of these metrics was based on the following considerations. Previous studies on cross-modal information integration (Rebai, Bernard, & Lannou, 1997; Qiu et al., 2006), including taste-visual integration (Xiao et al., 2014) suggested that presentation of incongruent stimuli is accompanied by an increase in the N430-620 amplitude which likely originates from the increased demand for cognitive resources required to resolve conflicting sensory information. Thus, the more contrasted are tasted and presented products, the higher amplitude of N430-620 is expected. Indeed, in our experimental data we observed a negative correlation between N430-620 amplitude and perceived product similarity. However, this metric may not be the optimal one to solve the task of identifying the most similar product because of a large inter-subject variability and low discriminative power between highly similar products.

Another considered metric was the power of induced gamma oscillations during 400-600 ms period after stimulus presentation. It is generally accepted that EEG synchronization in the gamma range is an important mechanism for generation of a multimodal object representation in our brain (Tallon-Baudry, & Bertrand, 2000; Yuval-Greenberg, & Deouell, 2007). Thus, we hypothesized that the more taste sensations correspond to the presented image, the greater the observed power of the induced gamma oscillations would be. And indeed, our experimental data revealed a strong positive correlation between this metric and perceived product similarity both at the individual and groups levels suggesting a universality of this approach. Furthermore, the overall results from EEG sessions well reflected the results from the hall-tests. Under “Open Eyes” conditions both hall-test and gamma power oscillations suggested that creamy MCCs were most similar to cereal bars, while under “Blind” condition – to waffles. At the same time, using gamma power oscillations allowed to identify granola as the most similar product to chocolate MCCs under “Open Eyes” condition, which was not possible based on hall-test only (granola, cereal bars and candies were considered equally similar). It was also suggested based on gamma power that the perceived similarity between chocolate MCCs and waffles may have been underestimated under “Blind” condition and, similar to creamy MCCs, they may be also associated with waffles under “Blind” condition testing.

## Conclusions

In this work we have suggested a novel approach to estimate perceived product similarity based on the power of induced gamma oscillations following presentation of congruent or incongruent visual stimuli. The results obtained using such an approach may be compared to traditional hall-tests but, unlike hall-tests, do not require large subject samples, do not pose any additional cognitive demands on respondents and do not depend on their language skills.

## References

Ariely, D., & Berns, G. S. (2010). Neuromarketing: the hope and hype of neuroimaging in business. Nature Reviews Neuroscience, 11(4), 284–292. doi: 10.1038/nrn2795.

Bassar-Eroğlu, C. et al. (1996). Gamma-band responses in the brain: a short review of psychophysiological correlates and functional significance. International Journal of Psychophysiology, 24 (1-2), 101–112. doi: 10.1016/S0167-8760(96)00051-7.

Qiu, J., Luo, Y. J., Wang, Q. H., Zhang, F. H., & Zhang, Q. L. (2006). Brain mechanism of Stroop interference effect in Chinese characters. Brain Res, 1072, 186–193. doi: 10.1016/j.brainres.2005.12.029.

Rebai, M., Bernard, C., & Lannou, J. (1997). The Stroop’s test evokes a negative brain potential. Neurosci, 91(1-2), 85–94. doi: 10.3109/00207459708986367.

Roininen, K. et al. (2001). Differences in health and taste attitudes and reported behaviour among Finnish, Dutch and British consumers: a cross-national validation of the Health and Taste Attitude Scales (HTAS). Appetite, 37(1), 33–45. doi: 10.1006/appe.2001.0414.

Stasi, A.. et al. (2018). Neuromarketing empirical approaches and food choice: A systematic review. Food Research International, 108, 650–664. doi: 10.1016/j.foodres.2017.11.049.

Tallon-Baudry, C., & Bertrand, O. (2000). Oscillatory gamma activity in humans and its role in object representation. International Journal of Psychophysiology, 38, 211–223. doi: 10.1016/s0167-8760(00)00166-5.

Vlăsceanu, S. (2014). Neuromarketing and evaluation of cognitive and emotional responses of consumers to marketing stimuli. Procedia: Social and Behavioral Sciences, 127, 753–757. doi: 10.1016/j.sbspro.2014.03.349.

We are what we eat: Healthy eating trends around the world (2015). Nielsen Global Health and Wellness Report. Retrieved from https://www.nielsen.com/wp-content/uploads/sites/3/2019/04/Nielsen20Global20Health20and20Wellness20Report.pdf.

What’s in our food and on our mind: Ingredient and dining-out trends around the world (2016). Nielsen Global Health and Wellness Report. Retrieved from https://www.nielsen.com/wp-content/uploads/sites/3/2019/04/global_ingredient_and_0ut_of_home_dining_trends_report.pdf.

Wright, K. B. (2005). Researching Internet-Based Populations: Advantages and Disadvantages of Online Survey Research, Online Questionnaire Authoring Software Packages, and Web Survey Services. Journal of Computer-Mediated Communication, 10(3). doi.org/10.1111/j.1083-6101.2005.tb00259.x.

Xiao, X. et al. (2014). The Taste-visual Cross-modal Stroop Effect: an Event related brain potential study. Neuroscience, 263, 250–256. doi: 10.1016/j.neuroscience.2014.01.002.

Yuval-Greenberg, S., & Deouell, L. Y. (2007). What You See Is Not (Always) What You Hear: Induced Gamma Band Responses Reflect Cross-Modal Interactions in Familiar Object Recognition. The Journal of Neuroscience, 27(5), 1090–1096. doi: 10.1523/JNEUROSCI.4828-06.2007.

